# Relationships between estimated autozygosity and complex traits in the UK Biobank

**DOI:** 10.1101/291872

**Authors:** Emma C Johnson, Luke M Evans, Matthew C Keller

## Abstract

Inbreeding increases the risk of certain Mendelian disorders in humans but may also reduce fitness through its effects on complex traits and diseases. Such inbreeding depression is thought to occur due to increased homozygosity at causal variants that are recessive with respect to fitness. Until recently it has been difficult to amass large enough sample sizes to investigate the effects of inbreeding depression on complex traits using genome-wide single nucleotide polymorphism (SNP) data in population-based samples. Further, it is difficult to infer causation in analyses that relate degree of inbreeding to complex traits because confounding variables (e.g., education) may influence both the likelihood for parents to outbreed and offspring trait values. The present study used runs of homozygosity in genome-wide SNP data in up to 400,000 individuals in the UK Biobank to estimate the proportion of the autosome that exists in autozygous tracts—stretches of the genome which are identical due to a shared common ancestor. After multiple testing corrections and controlling for possible sociodemographic confounders, we found significant relationships in the predicted direction between estimated autozygosity and three of the 26 traits we investigated: age at first sexual intercourse, fluid intelligence, and forced expiratory volume in 1 second. Our findings for fluid intelligence and forced expiratory volume corroborate those of several published studies while the finding for age at first sexual intercourse was novel. These results may suggest that these traits have been associated with Darwinian fitness over evolutionary time, although there are other possible explanations for these associations that cannot be eliminated. Some of the autozygosity-trait relationships were attenuated after controlling for background sociodemographic characteristics, suggesting that care needs to be taken in the design and interpretation of ROH studies in order to glean reliable information about the genetic architecture and evolutionary history of complex traits.

**Author Summary:** Inbreeding is well known to increase the risk of rare, monogenic diseases, and there has been some evidence that it also affects complex traits, such as cognition and educational attainment. However, difficulties can arise when inferring causation in these types of analyses because of the potential for confounding variables (e.g., socioeconomic status) to bias the observed relationships between distant inbreeding and complex traits. In this investigation, we used single-nucleotide polymorphism data in a very large (N > 400,000) sample of seemingly outbred individuals to quantify the degree to which distant inbreeding is associated with 26 complex traits. We found robust evidence that distant inbreeding is inversely associated with fluid intelligence and a measure of lung function, and is positively associated with age at first sex, while other trait associations with inbreeding were attenuated after controlling for background sociodemographic characteristics. Our findings are consistent with evolutionary predictions that fluid intelligence, lung function, and age at first sex have been under selection pressures over time; however, they also suggest that confounding variables must be accounted for in order to reliably interpret results from these types of analyses.

## Introduction

Inbreeding occurs when genetic relatives have offspring, and is associated with increased risk of disorders and decreased health and viability in offspring (1–3). This effect, called inbreeding depression, is thought to occur because natural selection more efficiently removes additive and dominant deleterious alleles, leaving the remaining deleterious alleles segregating in the population at a given time more recessive than otherwise expected (4), a phenomenon called directional dominance. Inbreeding is thought to be associated with lower fitness because it leads to long stretches of the genome that are autozygous—homozygous because the genomic segments inherited from each parent are from the same ancestor. Autozygosity reveals the full deleterious effects of recessive or partially recessive alleles that exist in these regions, and so individuals with increased autozygosity are more likely to exhibit deficits in traits that have been associated with Darwinian fitness over evolutionary time. Thus, one major reason for the interest in studying the effects of inbreeding on complex traits has been that such studies can provide insight into which traits have been under natural selection.

Because all humans are related to one another, even if distantly, inbreeding is a matter of degree. In the last decade, the increasing availability of genome-wide single nucleotide polymorphism (SNP) data has allowed scientists to infer degree of distant inbreeding, or the proportion of the genome that is autozygous, using runs of homozygosity (ROHs)—long stretches of SNPs that are homozygous (5). The total proportion of the genome contained within these homozygous regions is called *F*_*ROH*_ and has been shown to be the best genome-wide estimate of autozygosity (5,6). However, very large samples (e.g. *n* > 10,000) are required to detect likely effects of *F*_ROH_in outbred human populations because of the low variance in levels of genome-wide autozygosity in such populations. Previous studies of *F*_*ROH*_ in humans have found evidence consistent with inbreeding depression for several complex traits, including height, forced expiratory volume in one second (FEV1), educational attainment, and cognitive ability (*g*) (7–9), with less conclusive evidence for an effect of inbreeding on psychiatric disorders (10,11) or risk factors for late-onset diseases like hypertension and other cardiovascular disease (12,13). These observed associations with *F*_*ROH*_ may suggest that directional selection has acted on these traits ancestrally.

One challenge in autozygosity research in humans is in the causal interpretations of any observed *F*_ROH_-trait relationships. It is likely that propensity to outbreed (choosing mates who are genetically dissimilar) is related to multiple sociodemographic variables, such as education, religiosity, or socioeconomic status, that may also influence offspring trait values. In a recent study conducted in the Netherlands, a relatively small, densely populated country with a strong history of latitudinal religious assortment, Abdellaoui et al. (14) found a significant association between decreased *F*_*ROH*_ (i.e. less inbred) and increased risk for major depressive disorder (MDD); this counter-intuitive association disappeared when the models accounted for religious assortment. This suggests that the original *F*_*ROH*_ -MDD association occurred for sociological rather than genetic reasons: religious individuals had higher average levels of autozygosity than non-religious individuals, probably due to denominational restrictions on mate choice that were only recently relaxed (14), and religious individuals were less likely to experience MDD (15). In another recent study, the largest (N > 300,000) *F*_*ROH*_ analysis to date, Joshi et al. (2016) found a significant relationship between *F*_*ROH*_ and four complex traits: height, FEV1, cognitive ability (*g*), and educational attainment (7). When educational attainment was included as a covariate in the model as a proxy for SES, the effects for height,

FEV1, and cognitive ability remained significant. Because of the persistence of these effects after accounting for educational attainment, the authors conclude that the relationship they observed between *F*_*ROH*_ and the complex traits is likely a due to a genetic mechanism, directional dominance, rather than to sociodemographic confounds. However, the FROH-trait effect sizes decreased by ~20-35% after controlling for SES; it is possible that inclusion of additional, or more relevant, sociodemographic covariates could have changed these conclusions.

The findings from our work and others on the relationship between *F*_*ROH*_ and psychiatric disorders in ascertained samples (10,11,16–20) have been inconsistent and highlight concerns about the potential for unmeasured confounders to influence *F*_*ROH*_ results. Using the Psychiatric Genomics Consortium (PGC) MDD data from 9 samples, Power et al.(11) found a significant positive relationship between *F*_*ROH*_ and MDD in three German samples but, strangely, a significant negative relationship between *F*_*ROH*_ and MDD in six samples from non-German sites. Similarly, in 2012 we found a small but highly significant association between schizophrenia and *F*_*ROH*_ across 17 case-control datasets (total N = 21,844 (19)). However, in 2016 we published an independent replication using the same procedures as our previous study that found little to no evidence of an *F*_*ROH*_-schizophrenia association across 22 case-control datasets (total N = 39,830 (10)). We are uncertain how to explain these discrepancies, but we have hypothesized that unmeasured cofounding variables such as education, religiosity, and income can differentially bias such ROH findings across different sites, and that this problem is particularly salient in ascertained samples where cases and controls may be drawn from population that differ slightly on background sociodemographic characteristics. While such differences in ascertainment between cases and controls are unlikely to lead to significant allele frequency differences, and thus are unlikely to bias genome-wide association studies (GWAS), they could very easily lead to systematic case-control differences in *F*_*ROH*_, depending on the difference in degree of inbreeding in the populations from which cases and controls were drawn.

Here, we describe the most powerful investigation to date of the association of *F*_*ROH*_ with complex traits. We used whole-genome SNP and phenotypic data from the UK Biobank (total *n* ~ 100,000 - 400,000) to address two principal questions: (1) is there evidence consistent with directional dominance on traits related to fitness and health, such that increased *F*_*ROH*_ is associated with lower trait values? and (2) do F_R0H_-trait relationships persist after controlling for multiple background sociodemographic variables? This sample is population-based, reducing concerns about ascertainment-induced confounds, and includes information on multiple relevant sociodemographic control variables and traits previously associated with *F*_*ROH*_ (e.g. waist-to-hip ratio, grip strength, diastolic and systolic blood pressure, and fluid intelligence (7–9,11–13)), making it an ideal sample for investigating the relationship between distant inbreeding and complex traits.

## Methods

### Ethics Statement

This study utilized de-identified data from the UK Biobank. UK Biobank received ethical approval from the NHS National Research Ethics Service North West (11/NW/0382).

### UK Biobank Sample

Our study utilized data from up to 400,000 individuals (*n* varied by phenotype) with genotypes available from the UK Biobank, a population based sample from the United Kingdom. 502,682 individuals were recruited from 2006-2010 from 22 centers across the UK. Participants were given a touchscreen interview that included questions about demographic characteristics, health history, and lifestyle information (e.g. diet, alcohol intake, sleep habits), and some anthropometric and physical measures were collected. DNA was extracted from whole blood and genotyped using either the Affymetrix UK Biobank Axiom array or the Affymetrix UK BiLEVE Axiom array. Detailed genotyping and sample QC procedures are described in Bycroft et al.(21)

### Phenotypes

We examined 26 traits related to health, fitness, or sociodemographic characteristics (see Supplemental Materials for full description and field ID of individual measures). These included 17 continuous traits (age at first sexual intercourse, waist to hip ratio, height, body mass index (BMI), basal metabolic rate (BMR), diastolic and systolic blood pressure (BP), hand-grip strength (taking the maximum of left and right grip strength measurements), county-wide socioeconomic status (SES) as measured by the Townsend Deprivation Index (TDI), total household income (an ordinal variable of income brackets recoded to be numeric, ranging from 0 - 4), years of educational attainment (coded using ISCED classifications as in Okbay et al. (22)), fluid intelligence (FI), forced expiratory volume in 1 second (FEV1; a measure of lung functioning), FEV1 over forced vital capacity (FEV1/FVC), birth weight, neuroticism score, and body fat percentage) and 9 binary traits (ever smoked, ever drank alcohol, whether or not they were breastfed as a baby, whether or not they completed college, whether they specified participation in a religious group as a leisure activity, whether or not they had ever been diagnosed with diabetes, probable bipolar and/or major depression status, and whether or not they live in an urban or rural area). We excluded individuals who weighed less than 36.28 kg (~80 lbs), weighed more than 6.8 kg (~15 lbs) at birth, had systolic BP readings >200 mmHg or diastolic BP readings >120 mmHg, had a pulse <30 beats per minute or >130 beats per minute, were shorter than 120 cm (~3.93 ft), had a hip circumference <50 cm or >175 cm, had a waist circumference <40 cm or >160 cm, had grip strength >70 kg, or reported having had sex before 12 years of age. These exclusion criteria were chosen based on thresholds typically defined as being boundaries of normal physiological, anthropometric, or behavioral ranges and by checking for obvious outliers that may have been incorrect data entries. More information on specific phenotype derivations and calculations are included in the supplemental material. We standardized all quantitative phenotypes (within sex) before calculating their relationship with *F*_*ROH*_ for ease of comparison with Joshi et al.’s and others’ results (7).

### Quality Control (QC) and ROH calling Procedures

Because the sample was predominately European ancestry, we restricted analyses to individuals of European ancestry (n= 436,065) as identified by visual inspection of plots of genomic principle components. We followed sample and genotypic quality control that has become typical in ROH analyses. In particular, we excluded SNPs if they a) deviated from Hardy-Weinberg equilibrium at p<1×10^−6^, b) missingness proportion >0.02, or c) had a minor allele frequency (MAF) < 0.05. We also excluded individuals with a missing genotype call rate > 0.02, and we removed the minimum number of individuals so that all remaining subjects were unrelated at pihat > 0.2 (using GCTA’s --grm-cutoff option (23)) (n = 31,541 removed in total).

After QC, we pruned out SNPs that were in strong linkage disequilibrium with other SNPs by removing those that had a variance inflation factor > 10 (equivalent to an r^2^ of 0.90) between target SNPs and 50 surrounding SNPs (plink command: --indep 50 5 10). After these procedures, 263,609 SNPs and 404,524 individuals remained. For our main analysis, we called ROHs as being ≥65 homozygous SNPs in a row spanning at least 1000 kb, with no heterozygote calls and one missing variant call allowed, per recommendations from Howrigan et al. (2011) for genotype data of similar SNP density. We required ROHs to have a density greater than at least 1 SNP per 200 kb (the average density across the genome in the SNPs used in the analysis was 1 per 10 kb), and split an ROH into two if a gap >500 kb existed between consecutive homozygous SNPs. All analyses used the --homozyg commands in Plink 1.9 (24). After calling ROHs, we summed the total length of all autosomal ROHs for each individual and divided that by the total SNP-mappable distance (2.77×10^9^ bases) to calculate *F*_*ROH*_, the proportion of the genome likely to be autozygous. In addition to calling ROHs, we also calculated a measure of SNP-by-SNP homozygosity (*F*_*SNP*_,) for each individual, using the --het flag in Plink 1.9 (24):

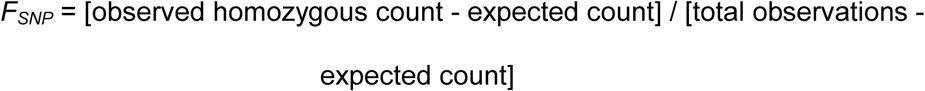

Because it is calculated with genotyped SNPs, *F*_*SNP*_, is a measure of excess homozygosity at common SNPs.

### ROH Burden Analysis

*F*_*roh*_ was used as the primary predictor of the traits of interest in analyses described below. The distributions of ROH lengths and *F*_*ROH*_ are shown in Figure S1 (see Supplement). We regressed each trait (Y) on *F*_*ROH*_ using the model in the equation below, where 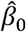 is the intercept, *C* is a matrix of covariates (including e.g. the first 20 principle components) and ε represents the residual error term.

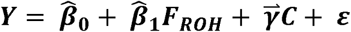

As noted above, all quantitative phenotypes were standardized to intra-sex 0 for ease of comparison with previous findings in the literature. In addition, for ease of interpretation, we reverse-coded some of the phenotypes such that lower values represented what we thought were likely to be lower fitness and/or less desirable outcomes (e.g. disease diagnosis was coded as ‘0’ while no diagnosis as ‘1’, and TDI was reverse-coded such that lower values represented greater material poverty). We were primarily interested in the estimate of 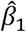 which represents the association of *F*_*ROH*_ with the trait, controlling for covariates (although in one set of models, described below, we were also interested in the effect of *F*_*SNP*_, on the trait). For binary traits, we ran logistic regression models with the same covariates as in the linear regression models for quantitative traits.

We ran a total of three sets of models for each trait. The first set of models was designed to test for a simple relationship between *F*_*ROH*_ and the traits listed above. Because confounding factors such as population stratification, SNP missingness, call quality, and plate effects can influence *F*_*ROH*_, we included the batch number, percentage of missing SNP calls per sample in the raw data, and the first 20 ancestry principal components (calculated within individuals of European ancestry), as well as age, age^2^, and sex, in all of the regression models unless explicitly stated.

In our second set of models, we tested whether background sociodemographic characteristics mediated F_ROH_-trait relationships. In addition to the above covariates, in these models we also included income, years of educational attainment, Townsend Deprivation Index (a measure of the amount of material deprivation in a given region), and whether subjects attended college, lived in an urban area, participated in a religious group as a leisure activity, and reported being breastfed as an infant. Although the covariates of true interest are those measured on the parents (whose sociodemographic traits may influence mate choice), parental information was unavailable (other than breastfeeding, which is associated with mother’s socioeconomic status (25)), and so we used the subjects’ own values on these traits as the best available proxies for characteristics of the their parents. To formally test whether the seven sociodemographic variables, in combination, significantly mediated the *F*_*ROH*_ associations identified in the first model, we followed Kenny and Judd’s recommendations (26) for calculating the indirect effect size and then bootstrapped with 1,000 resamples to get the 99% confidence intervals around the indirect path coefficients for any significant *F*_ROH_-trait associations we observed in our first set of models.

In our third set of models, we tested the degree to which observed F_ROH_-trait relationships were due to homozygosity at common versus rare alleles. To do this, we included *F*_*SNP*_, as a covariate in addition to the covariates from the second set of models above. Because common SNPs can often predict (are in linkage disequilibrium with) other common SNPs but typically poorly predict rare SNPs, *F*_*SNP*_, captures effects of homozygosity at common SNPs only whereas *F*_*ROH*_ captures the effects of homozygosity at both common and rare SNPs (5). In the Supplement (Table S3), we demonstrate via simulation that entering both *F*_*SNP*_, and *F*_*ROH*_ as predictors simultaneously in the regression equation allows insight into the degree to which observed inbreeding effects are due to homozygosity at common versus rare alleles.

## Results

The distribution of ROH lengths, *F*_*ROH*_, and *F*_*SNP*_, are shown in Figures S1-S2, and descriptive statistics are given in Table S1. Using a Bonferroni correction based on testing 26 traits (alpha = .002), we observed significant negative associations between *F*_*ROH*_ and grip strength, height, fluid intelligence score (FI), and forced expiratory volume in one second (FEV1), and observed significant positive associations between *F*_*ROH*_ and age at first sexual intercourse (AFS) and religious group participation (Table 1 and Figure 1). The associations we found between *F*_*ROH*_ and FI, FEV1, and height replicate three of Joshi et al.’s four significant findings. To our surprise, we did not replicate their significant relationship between *F*_*ROH*_ and educational attainment. As a post hoc analysis, we also tested the relative importance of recent vs. distant inbreeding by calculating *F*_*ROH*_ from longer ROHs (indicative of closer inbreeding) and comparing to the effect of *F*_*ROH*_ from shorter ROHs (a proxy for more distant inbreeding). We defined recent inbreeding as the proportion of the genome contained in autozygous regions longer than 8.5 Mb (*F_ROH,recent_*) and distant inbreeding as the proportion of the genome in autozygous regions shorter than 8.5 Mb (*F*_*ROH,distant*_), as *F*_*ROH,recent*_ and *F*_*ROH,distant*_ had approximately equal variances (4.5e-6 and 4.3e-6, respectively) in our sample. An autozygous segment spanning < 8.5 Mb should originate from a common ancestor at least 6 generations ago on average (27). Results for more recent inbreeding were similar to the full *F*_*ROH*_ models: income, grip strength, height, fluid intelligence score (FI), forced expiratory volume in one second (FEV1), age at first sexual intercourse (AFS), and religious group participation were all associated with *F_ROH.recent_,* with the same direction of effect as the original models. Similarly, AFS, FEV1, FI, religious group attendance, and ever drink (such that being more autozygous was associated with a lower likelihood of having ever drank alcohol) were significantly associated with *F*_*ROH,distant*_, while its associations with income, grip strength, and height were not (Table S4).

**Table 1.**
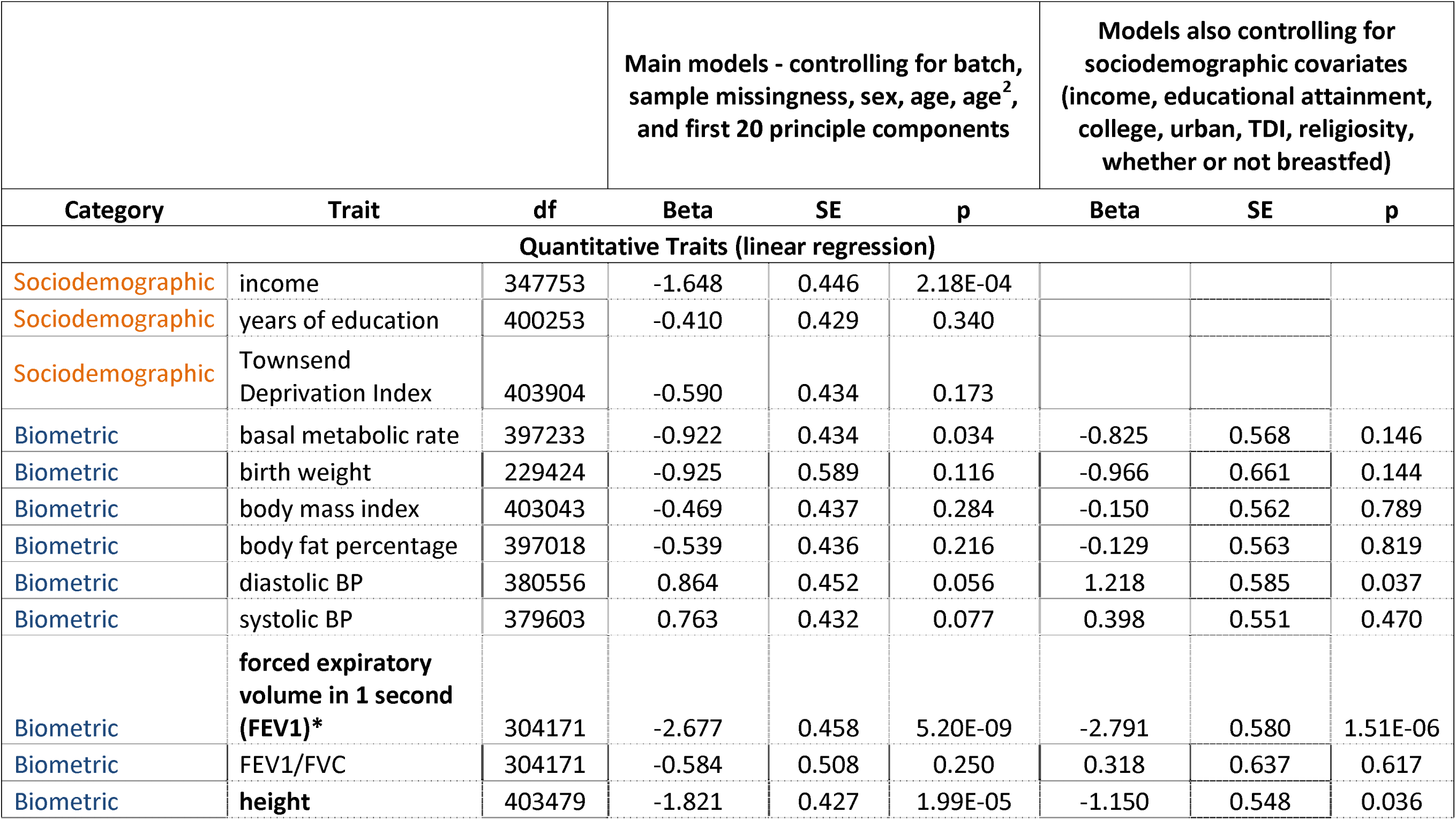
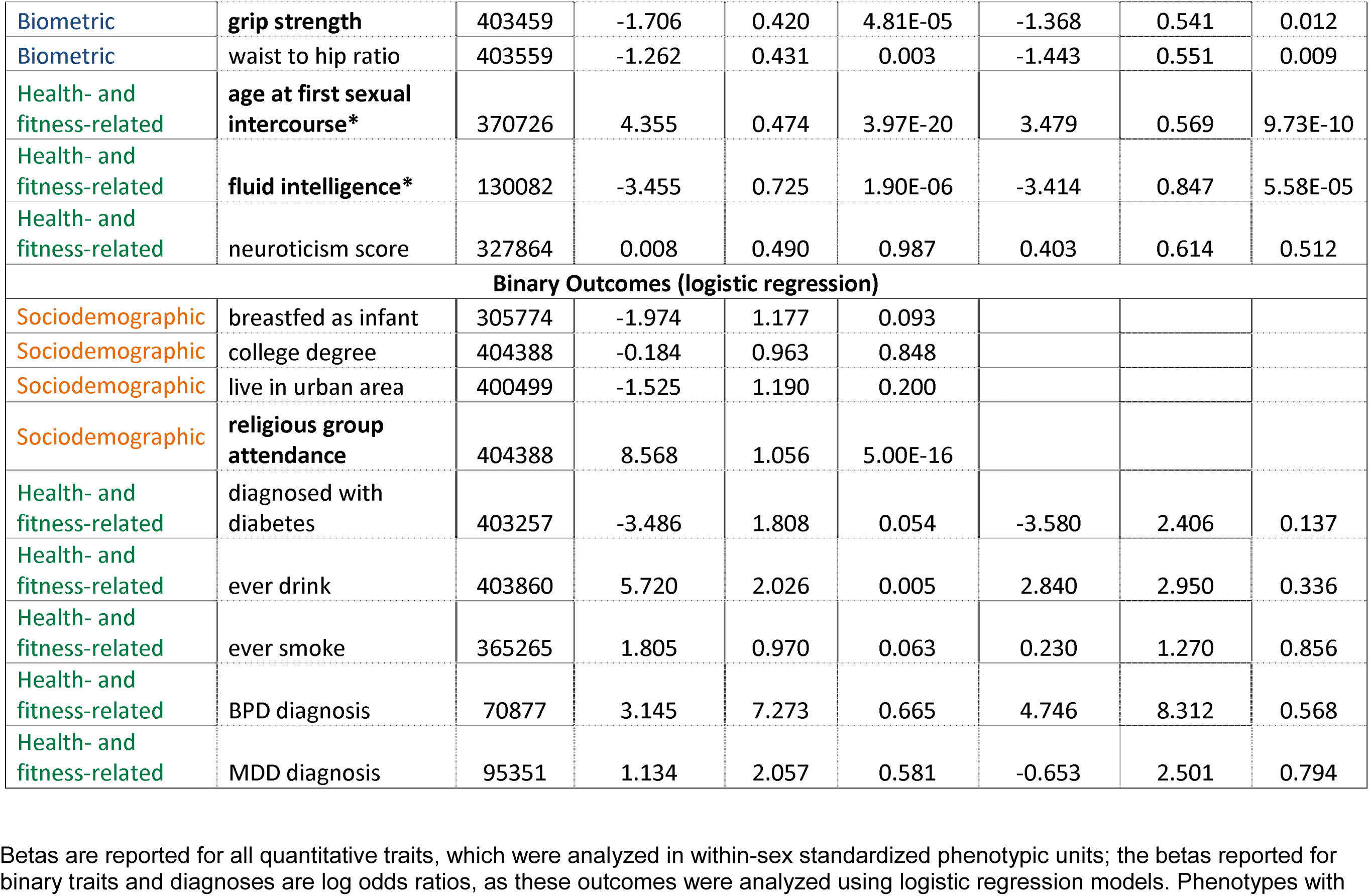
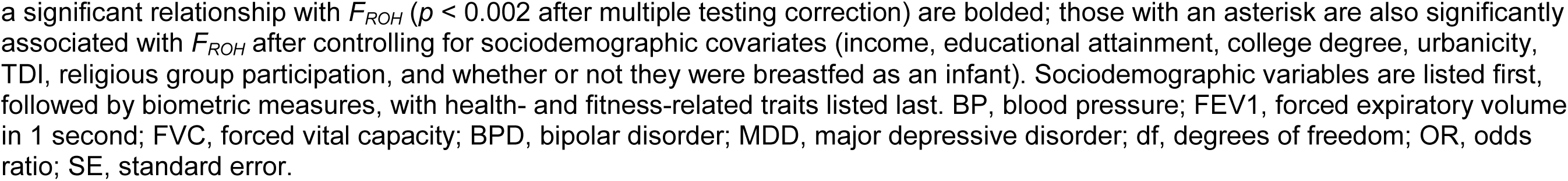
Association of *F*_*ROH*_ with 26 traits, in two sets of models: 1) controlling for age, age^2^, sex, the first 20 principal components, sample missingness, and batch number as covariates, and 2) also controlling for sociodemographic variables.

|  |  |  | Main models - controlling for batch, sample missingness, sex, age, age<sup>2</sup>, and first 20 principle components |  |  | Models also controlling for sociodemographic covariates (income, educational attainment, college, urban, TDI, religiosity, whether or not breastfed) |  |  |
| --- | --- | --- | --- | --- | --- | --- | --- | --- |
| Category | Trait | df | Beta | SE | p | Beta | SE | p |
| Quantitative Traits (linear regression) |  |  |  |  |  |  |  |  |
| Sociodemographic | income | 347753 | -1.648 | 0.446 | 2.18E-04 |  |  |  |
| Sociodemographic | years of education | 400253 | -0.410 | 0.429 | 0.340 |  |  |  |
| Sociodemographic | Townsend Deprivation Index | 403904 | -0.590 | 0.434 | 0.173 |  |  |  |
| Biometric | basal metabolic rate | 397233 | -0.922 | 0.434 | 0.034 | -0.825 | 0.568 | 0.146 |
| Biometric | birth weight | 229424 | -0.925 | 0.589 | 0.116 | -0.966 | 0.661 | 0.144 |
| Biometric | body mass index | 403043 | -0.469 | 0.437 | 0.284 | -0.150 | 0.562 | 0.789 |
| Biometric | body fat percentage | 397018 | -0.539 | 0.436 | 0.216 | -0.129 | 0.563 | 0.819 |
| Biometric | diastolic BP | 380556 | 0.864 | 0.452 | 0.056 | 1.218 | 0.585 | 0.037 |
| Biometric | systolic BP | 379603 | 0.763 | 0.432 | 0.077 | 0.398 | 0.551 | 0.470 |
| Biometric | forced expiratory volume in 1 second (FEV1)* | 304171 | -2.677 | 0.458 | 5.20E-09 | -2.791 | 0.580 | 1.51E-06 |
| Biometric | FEV1/FVC | 304171 | -0.584 | 0.508 | 0.250 | 0.318 | 0.637 | 0.617 |
| Biometric | height | 403479 | -1.821 | 0.427 | 1.99E-05 | -1.150 | 0.548 | 0.036 |
| Biometric | **grip strength** | 403459 | -1.706 | 0.420 | 4.81E-05 | -1.368 | 0.541 | 0.012 |
| Biometric | waist to hip ratio | 403559 | -1.262 | 0.431 | 0.003 | -1.443 | 0.551 | 0.009 |
| Health- and fitness-related | **age at first sexual intercourse\*** | 370726 | 4.355 | 0.474 | 3.97E-20 | 3.479 | 0.569 | 9.73E-10 |
| Health- and fitness-related | **fluid intelligence\*** | 130082 | -3.455 | 0.725 | 1.90E-06 | -3.414 | 0.847 | 5.58E-05 |
| Health- and fitness-related | neuroticism score | 327864 | 0.008 | 0.490 | 0.987 | 0.403 | 0.614 | 0.512 |
| **Binary Outcomes (logistic regression)** |  |  |  |  |  |  |  |  |
| Sociodemographic | breastfed as infant | 305774 | -1.974 | 1.177 | 0.093 |  |  |  |
| Sociodemographic | college degree | 404388 | -0.184 | 0.963 | 0.848 |  |  |  |
| Sociodemographic | live in urban area | 400499 | -1.525 | 1.190 | 0.200 |  |  |  |
| Sociodemographic | **religious group attendance** | 404388 | 8.568 | 1.056 | 5.00E-16 |  |  |  |
| Health- and fitness-related | diagnosed with diabetes | 403257 | -3.486 | 1.808 | 0.054 | -3.580 | 2.406 | 0.137 |
| Health- and fitness-related | ever drink | 403860 | 5.720 | 2.026 | 0.005 | 2.840 | 2.950 | 0.336 |
| Health- and fitness-related | ever smoke | 365265 | 1.805 | 0.970 | 0.063 | 0.230 | 1.270 | 0.856 |
| Health- and fitness-related | BPD diagnosis | 70877 | 3.145 | 7.273 | 0.665 | 4.746 | 8.312 | 0.568 |
| Health- and fitness-related | MDD diagnosis | 95351 | 1.134 | 2.057 | 0.581 | -0.653 | 2.501 | 0.794 |

Betas are reported for all quantitative traits, which were analyzed in within-sex standardized phenotypic units; the betas reported for binary traits and diagnoses are log odds ratios, as these outcomes were analyzed using logistic regression models. Phenotypes with
a significant relationship with $F_{ROH}$ ( $p < 0.002$ after multiple testing correction) are bolded; those with an asterisk are also significantly associated with $F_{ROH}$ after controlling for sociodemographic covariates (income, educational attainment, college degree, urbanicity, TDI, religious group participation, and whether or not they were breastfed as an infant). Sociodemographic variables are listed first, followed by biometric measures, with health- and fitness-related traits listed last. BP, blood pressure; FEV1, forced expiratory volume in 1 second; FVC, forced vital capacity; BPD, bipolar disorder; MDD, major depressive disorder; df, degrees of freedom; OR, odds ratio; SE, standard error.

**Figure 1.**
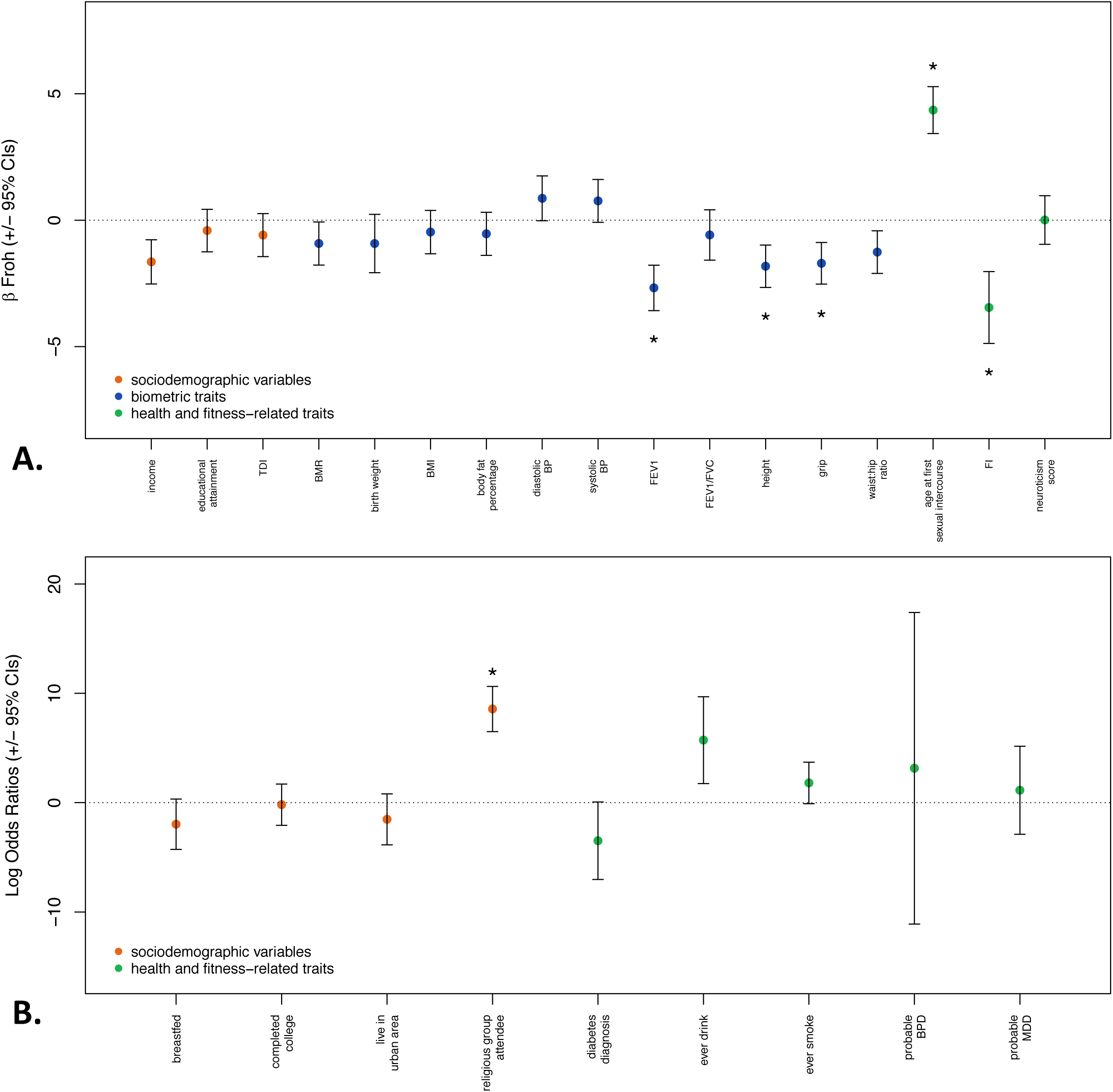
Beta *F*_*ROH*_ and 95% confidence intervals from main regression models controlling for minimal covariates (20 ancestry principal components, genotype batch, per-sample SNP missingness, age, age^2^, and sex). Significant estimates (at p < 0.002 - corrected for multiple testing) are starred (religious group attendance as a leisure activity, age at first sexual intercourse, FEV1, FI, height, and grip strength). **A.** All quantitative traits were analyzed in intra-sex standardized phenotypic units in linear regression models. **B.** Binary traits and diagnoses were analyzed using logistic regression models (the log odds ratios are reported). BMI, body mass index; BMR, basal metabolic rate; BPD, bipolar disorder; MDD, major depression; CI, confidence interval; FI, fluid intelligence; FEV1, forced expiratory volume in 1 second; TDI, Townsend Deprivation Index.

When we included the seven sociodemographic variables as covariates in the regression models (other than those predicting sociodemographic variables), the betas associated with *F*_*ROH*_ decreased for AFS, grip strength, height, and FI (by 20.1%, 19.8%, 36.8%, and 1.2%, respectively) and increased for FEV1 (by 4.2%) (see Table 1, Figure 2). AFS, FI, and FEV1 remained significantly associated with *F*_*ROH*_ whereas the associations with height and grip strength became non-significant; no significant indirect mediation effect of the sociodemographic variables in combination was found (see Supplemental Materials for further discussion). Furthermore, the association between *F*_*ROH,distant*_ and ever drink disappeared after controlling for the sociodemographic covariates, as did the associations between *F*_*ROH,recent*_ and grip strength, height, and FI (Table S5). Finally, we tested whether the effect of *F*_*ROH*_differed by sex by including sex* *F*_*ROH*_ interaction terms in each of the second set of models, but observed no significant sex-by-F_ROH_ interactions for any of the traits.

**Figure 2.**
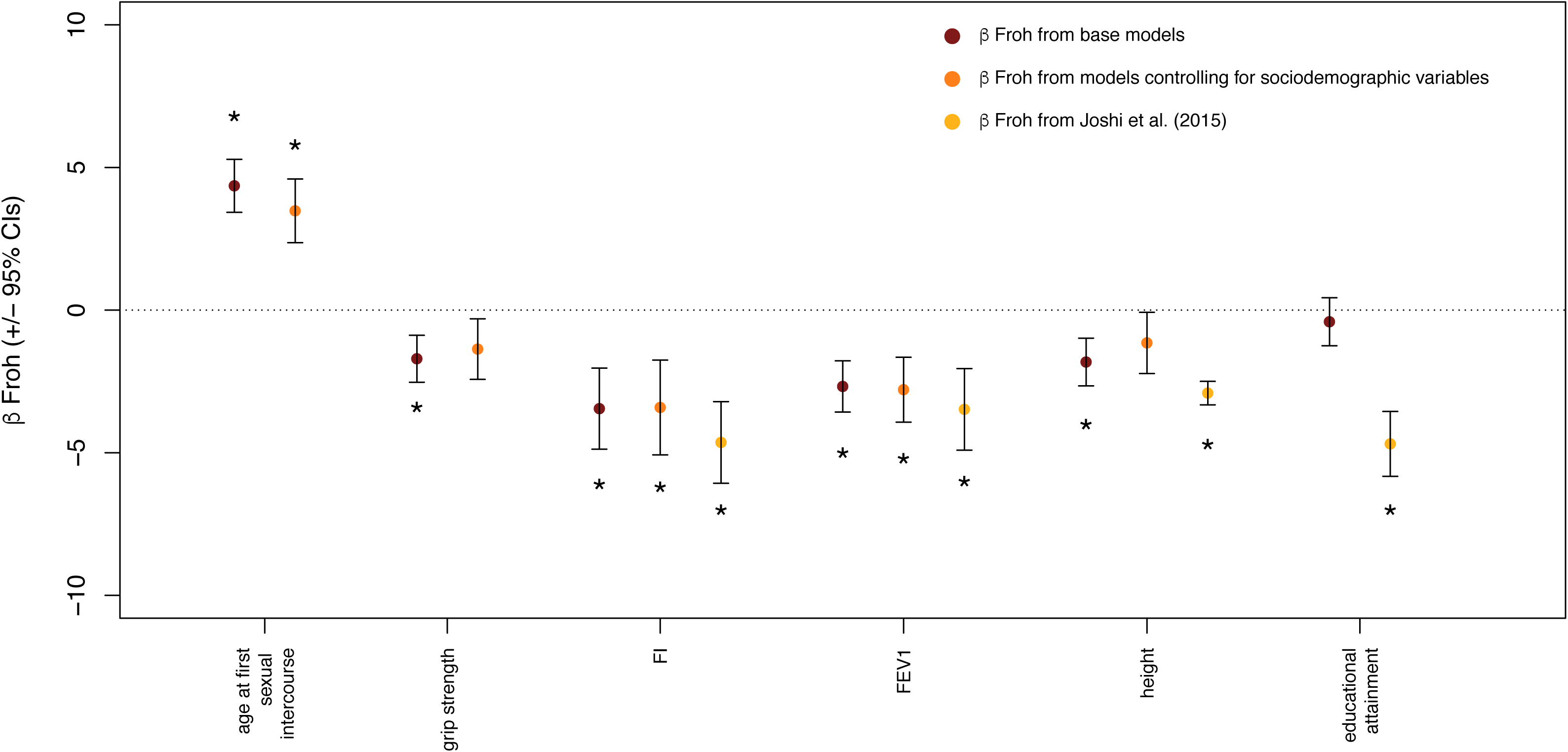
Comparison with estimates from Joshi et al. 2015, and some evidence that sociodemographic background variables attenuate the relationship between *F*_*ROH*_ and complex traits. Plot shows the Beta *F*_*ROH*_ and 95% confidence interval in within-sex standardized phenotypic units for the five quantitative traits that were significantly associated with *F*_*ROH*_ in the main models (Fig 1), as well as educational attainment, which was significantly associated with autozygosity in Joshi et al.’s study(7). Estimates that were statistically significant after multiple testing corrections are starred for each set of models. After controlling for background sociodemographic characteristics, age at first sexual intercourse, FEV1, and FI were still statistically significant in our study. The effect sizes for AFS, grip strength, FI, and height all decreased after controlling for sociodemographic variables. The effect sizes from our analyses were smaller for all four of the phenotypes also measured in Joshi et al.’s study. FI, fluid intelligence; FEV1, forced expiratory volume in 1 second; CI, confidence interval.

In our final set of models, where excess SNP-by-SNP homozygosity (*F*_*SNP*_,) was included as an additional covariate, AFS and FI remained significantly associated with *F*_*ROH*_ after accounting for multiple testing and FEV1 was marginally significant (Table 2). Waist-to-hip ratio was significantly associated with *F*_*SNP*_, but not *F*_*ROH*_, suggesting that higher homozygosity at common but not rare variants is related to increased waist-to-hip ratio.

**Table 2.** Effects of both *F*_*ROH*_and excess SNP-by-SNP homozygosity, measured by *F*_*SNP*_, controlling for the covariates in the previous models (age, sex, batch number, per-sample SNP missingness, the first 20 principal components, and background sociodemographic variables.)

| Model controlling for $F_{SNP}$, batch, sample missingness, sex, age, age<sup>2</sup>, and first 20 principal components | | | | | | | |
| --- | --- | --- | --- | --- | --- | --- | --- |
| Category | Trait | Beta<sub>FROH</sub> | SE<sub>FROH</sub> | p<sub>FROH</sub> | Beta<sub>FSNP</sub> | SE<sub>FSNP</sub> | p<sub>FSNP</sub> |
| Quantitative Traits (linear regression) |  |  |  |  |  |  |  |
| biometric | basal metabolic rate | -0.883 | 0.678 | 0.193 | 0.056 | 0.361 | 0.876 |
| biometric | birth weight | -1.443 | 0.792 | 0.069 | 0.465 | 0.426 | 0.275 |
| biometric | body mass index | -0.230 | 0.672 | 0.732 | 0.078 | 0.358 | 0.828 |
| biometric | body fat percentage | -0.353 | 0.672 | 0.599 | 0.219 | 0.357 | 0.540 |
| biometric | diastolic BP | 1.334 | 0.700 | 0.057 | -0.113 | 0.373 | 0.762 |
| biometric | systolic BP | 0.725 | 0.658 | 0.271 | -0.318 | 0.351 | 0.365 |
| biometric | forced expiratory volume in 1 second (FEV1) | -2.010 | 0.688 | 0.0035 | -0.762 | 0.361 | 0.035 |
| biometric | FEV1/FVC | -0.110 | 0.755 | 0.884 | 0.418 | 0.396 | 0.291 |
| biometric | height | -0.950 | 0.655 | 0.147 | -0.195 | 0.349 | 0.576 |
| biometric | grip strength | -0.620 | 0.645 | 0.337 | -0.728 | 0.342 | 0.033 |
| biometric | waist to hip ratio | -0.137 | 0.658 | 0.835 | -1.273 | 0.351 | 2.88E-04 |
| health- and fitness-related | age at first sexual intercourse | 2.858 | 0.676 | 2.35E-05 | 0.602 | 0.354 | 0.089 |
| health- and fitness-related | **fluid intelligence** | -3.238 | 1.016 | 0.001 | -0.171 | 0.544 | 0.753 |
| health- and fitness-related | neuroticism score | 0.432 | 0.732 | 0.555 | -0.029 | 0.389 | 0.941 |
| **Binary Outcomes (logistic regression)** |  |  |  |  |  |  |  |
| health- and fitness-related | diagnosed with diabetes | -3.840 | 3.051 | 0.208 | 0.252 | 1.812 | 0.890 |
| health- and fitness-related | ever drink | 2.304 | 3.749 | 0.539 | 0.520 | 2.241 | 0.817 |
| health- and fitness-related | ever smoke | -1.699 | 1.510 | 0.261 | 1.881 | 0.796 | 0.018 |
| health- and fitness-related | BPD diagnosis | 7.619 | 10.645 | 0.474 | -2.864 | 6.632 | 0.666 |
| health- and fitness-related | MDD diagnosis | -0.877 | 3.028 | 0.772 | 0.220 | 1.679 | 0.896 |

Phenotypes with a significant relationship with $F_{ROH}$ ( $p < 0.003$ after multiple testing correction) are bolded, while those with a significant relationship with $F_{SNP}$ are italicized. The quantitative traits (analyzed via linear regression) are listed first in the table, followed by diagnoses and binary traits. BP, blood pressure; FEV1, forced expiratory volume in 1 second; FVC, forced vital capacity; BPD, bipolar disorder; MDD, major depressive disorder; df, degrees of freedom; SE, standard error.

## Discussion

### Overview of findings

We replicated several previous associations between *F*_*ROH*_ and fitness-related traits, identified a novel association between *F*_*ROH*_ and a reproductive phenotype (age at first sexual intercourse), and found weak evidence that background sociodemographic characteristics may be partially mediating a few of the observed relationships between *F*_*ROH*_ and complex traits (Figure 2). In particular, we found robust evidence that fluid intelligence (FI), forced expiratory volume in one second (FEV1), and age at first sexual intercourse (AFS) are associated with *F*_*ROH*_ (Tables 1 and 2), while grip strength and height’s relationships with *F*_*ROH*_ were attenuated enough to become non-significant after controlling for background sociodemographic variables. The strength of *F*_*ROH*_ associations for more recent inbreeding was similar or stronger than those for more distant inbreeding, except, interestingly, for participation in a religious group. When we accounted for SNP-by-SNP homozygosity in the model, AFS and FI were still significantly associated with *F*_*ROH*_, consistent with their relationships with *F*_*ROH*_ being more strongly driven by homozygosity at rare rather than common variants. Certain other associations were likely due to social rather than genetic causes; for example, it is much more plausible that non-religious individuals tend to outbreed at higher rates and have less religious offspring than that autozygosity causes individuals to be more religious.

### Comparison with previous results

Our results largely agree with recent reports (7–9) on the relationships between estimated autozygosity and complex traits in population-based samples. Replicating both Howrigan et al. (8) and Joshi et al. (7) as well as previous pedigree studies (28), we found a significant, negative relationship between *F*_*ROH*_ and fluid intelligence. In addition, we replicated Joshi et al.’s (7) finding of a significant relationship between increased *F*_*ROH*_ and decreased FEV1. We initially observed a significant association between increased *F*_*ROH*_ and decreased height, as did Joshi et al.(7) and Verweij et al.(9), but this association was attenuated in our sample after controlling for background sociodemographic variables and did not meet statistical significance after Bonferroni corrections (Table 1, Figure 2). Our initial results (Table 1) were consistent with previous findings for an effect of inbreeding depression on grip strength (9), though this association appears to be largely due to homozygosity at common rather than rare variants (Table 2).

Despite the general consistency across reports on *F*_*ROH*_-complex trait associations, there were two differences between our results and those from earlier studies. First, educational attainment (in years of education) was not significantly associated with *F*_*ROH*_ in any of our models, contrary to previous reports (7,9). We found a suggestive (*p* = 2.18e-4) relationship between *F*_*ROH*_ and income (which itself was correlated with years of education at *r* = 0.37), but we found no evidence for an association between *F*_*ROH*_ and either years of education or the binary variable measuring whether or not an individual attended college. The reason for the discrepancy in findings for education is unlikely to be due to sampling variability because the two confidence intervals do not overlap (Figure 2). One possibility is that educational attainment is less correlated with geographic mobility (and the tendency to outbreed) in the UK compared to other countries previously investigated, and Joshi et al. (7) did report significant heterogeneity of the *F*_*ROH*_-education association across sites. Moreover, of the 5 cohorts from the UK investigated by Joshi et al. (7), two (GRAPHIC and LBC1936) showed associations in the opposite direction of the overall association (see their Extended Data Figure 2). Thus, it is possible that the *F*_*ROH*_-educational1 attainment relationship might be different in the UK than is typical in other societies. Furthermore, the association we found between height and autozygosity was attenuated (by ~37%) when we accounted for sociodemographic covariates, and was somewhat smaller than that found by previous studies even when we did not control for sociodemographic variables (e.g. a 1% increase in *F*_*ROH*_ predicted a decrease of ~.03 s.d. of height in previous studies (7,29) versus a decrease of ~.02 s.d. in the current study). Nevertheless, the confidence intervals for Joshi et al.’s (2015) and our observed association between height and *F*_*ROH*_ overlapped (Figure 2), suggesting that sampling variability could be a reason for the discrepancy in height findings.

In comparing results across recent publications and the current one, it is important to note the differences in populations, samples, and measurements across studies. Both Howrigan et al. (8) and Joshi et al. (7) took a meta-analytic approach, conducting *F*_*ROH*_ analyses in each contributing sample separately, and then combining across samples, controlling for relevant covariates (e.g. dataset, country of data collection). Joshi et al. in particular analyzed a much more diverse overall sample than the present study, which included multiple cohorts from European, African, and Asian populations. Another difference is in the measurement of intelligence across studies: our measurement for general cognitive ability was the unweighted sum of the number of 13 fluid intelligence questions answered correctly, given as part of the UK Biobank’s cognitive function assessment, while Howrigan et al. (8) converted the scores from each contributing sample’s measure of general cognitive ability (e.g. WAIS-R, Cattell Culture Fair Test) into z-scores (to avoid bias from different measurement schemes across samples), and Joshi et al. used *g* as their measure of general cognitive ability, “calculated as the first unrotated principal component of test scores across diverse domains of cognition”. Finally, our regression models controlled for the first 20 ancestry principal components, while Howrigan et al. controlled for the first 10 and Joshi et al. the first 3.

### Possible evolutionary interpretations

There are two major evolutionary theories for why inbreeding depression occurs (4): the overdominance hypothesis posits that an overall loss of heterozygosity at loci governed by heterozygote advantage leads to inbreeding depression, while the partial dominance theory postulates that inbreeding depression occurs as selection acts most efficiently on the most additive and dominant deleterious mutations, purging those from the population while leaving behind the more rare, partially recessive deleterious alleles. This second hypothesis, partial dominance, is widely accepted as the more likely mechanism of inbreeding depression (3,30). The robust associations we observed between *F*_*ROH*_ and AFS, FI, and FEV1, even after controlling for homozygosity at common variants with *F*_*SNP*_, suggest that the variants contributing to lower trait values are biased toward being rare and recessive, consistent with predictions from a partial dominance model of inbreeding depression (5) and consistent with the hypothesis that these traits, or traits genetically correlated with them, have been under directional selection over evolutionary time. Cognitive ability, including intelligence test scores, is a predictor of multiple Darwinian fitness-related outcomes, including overall health and lifespan (8,31). FEV1 is correlated with mortality and lifespan (32–35), traits that are components of fitness and thus more likely to have been under directional selection over evolutionary history (36). Thus, our replication of the associations between autozygosity and FEV1 and FI adds to a body of evidence that these traits, or traits genetically correlated with them, have been under directional selection over evolutionary history, leading to deleterious variants that are biased toward being rarer and more recessive than otherwise expected.

The positive relationship we observed between AFS and *F*_*ROH*_ is a novel finding, to the best of our knowledge. The *F*_*ROH*_-AFS association remained statistically significant after controlling for sociodemographic variables and homozygosity at common variants (*F*_*SNP*_,). Although novel, the finding is consistent with a body of research suggesting that reproductive traits, like AFS, in non-human populations are under more intense selection pressures than non-fitness traits (5,37)). If autozygosity causally influences AFS (see “Limitations” below), there are two possible evolutionary interpretations. First, it is possible that early sex itself was advantageous in ancient human history due to a prolonged reproductive period. A second possibility is that the observed association between autozygosity and AFS is due to selection on a genetically correlated trait, such as sexual attractiveness (38,39).

### Limitations

There were two central limitations in the current study. The most important one, which applies equally to all other *F*_*ROH*_ studies that we are aware of, is that ROH associations might be due to third-variable explanations. Unlike GWAS analyses, where parental or offspring sociodemographic traits are unlikely to be associated with allele frequencies and therefore are unlikely to bias GWAS results, it takes only a single generation of parental inbreeding to strongly influence *F*_*ROH*_ levels in offspring. For example, higher income might be associated with greater opportunities to meet mates of diverse origins and to higher outbreeding; offspring of higher income parents might thereby have not only lower levels of autozygosity, on average, but might also differ on any traits influenced genetically or environmentally by parental income. While sociodemographic confounding is particularly problematic in ascertained samples where cases and controls are drawn from different populations (e.g. cases drawn from a psychiatric hospital, controls from a nearby university), the possibility of confounding cannot be eliminated, even in population-based samples, unless relevant sociodemographic variables among parents are measured and controlled for or other (e.g., within-family) designs are used. For example, in a study of approximately 2,000 individuals of Dutch ancestry, Abdellaoui et al.(40) found only a weak association between *F*_*ROH*_ and the subjects’ own educational attainment (*p* = 0.045), but found highly significant negative associations between the subject’s *F*_*ROH*_ and their parents’ educational attainment (*p*_*father*_ < 10^−5^, *P*_*mother*_ = 9e^−5^). These relationships were entirely mediated by the geographic distance between parents’ birthplaces, such that parents with higher educational attainment tended to be more geographically mobile, increasing their chances of mating with someone genetically dissimilar from themselves and thus creating systematic differences in levels of inbreeding across levels of educational attainment in their offspring.

Having information on parents’ birth location, education, income, mobility, level of religious involvement, and so forth is important in order to control for the possibility that these sociodemographic variables are associated with both higher levels of (distant) inbreeding and lower offspring trait values. Unfortunately, the UK Biobank has limited parental information other than indirect measures such as whether one was breastfed. In the current study, we used sociodemographic responses of the offspring as imperfect proxies for parental responses, which is effective only to the degree that offspring values on these sociodemographic variables are positively correlated with their parents’ values. For example, parental educational (*r* = 0.25 - 0.40; (41,42)), income (r = .60; (42)), and religiosity (43) are imperfectly correlated between parents and offspring in Great Britain. These imperfect correlations imply that the true mediating influences of the sociodemographic variables on observed *F*_*ROH*_-trait relationships were likely to be underestimated in the present report, and thus causal interpretation of our results may not be warranted.

The second limitation to the current study is that we did not have access to all of the phenotypes studied in recent articles such as Verweij et al. (9) or Joshi et al. (7) (e.g. the cholesterol measures in Joshi et al.), so we could not attempt to fully replicate these previous investigations.

### Summary

We found several significant associations between estimated autozygosity and sociodemographic, anthropometric, health, and otherwise fitness-related traits, including whether or not a person attends a religious group as a leisure activity, AFS, grip strength, height, FI, and FEV1. All effects were in the direction that would be predicted by evolutionary hypotheses (i.e. higher inbreeding associated with lower fitness). When controlling for measures of background sociodemographic characteristics (educational attainment, college education, income, urbanicity, TDI, religious participation, and whether or not an individual was breastfed) - which should at least partially reflect parental characteristics - we found that two (height and grip strength) of the five significant *F*_*ROH*_-trait associations were attenuated and became non-significant, while AFS, FI, and FEV1 remained significantly associated with *F_RoH_.* The fact that the associations between estimated autozygosity and both grip strength and height were reduced after controlling for the additional covariates suggests that these relationships might not hold up if relevant confounder variables in parents had been controlled for, and we cannot eliminate the possibility that the other *F*_*ROH*_-trait associations we report here would not also be attenuated or eliminated in this situation.

Nevertheless, our results are consistent with the hypothesis that natural selection has biased the alleles contributing to lower fluid intelligence, later age at first sex, and poorer lung functioning (as measured by FEV1) toward being rare and recessive. These findings generally replicate previous findings in humans (7–9), and are consistent with similar ones from non-human populations (37,44,45). This cumulative evidence may well reflect the detrimental effects of autozygosity on complex traits, revealing ancient selection pressures on these or correlated traits. However, the fact remains that even in very large, well-powered, unascertained samples such as this one, it is exceedingly difficult to make definitive statements about the underlying causal mechanism of observed relationships between *F*_*ROH*_ and complex traits.

## Acknowledgements

We thank Chick Judd for his statistical guidance on testing mediation models, as well as Brooke Huibregtse for her valuable input regarding some of the UK Biobank phenotypes. This work was supported by R01 MH100141 (MCK). This research has been conducted using the UK Biobank Resource under Application Numbers ‘1665’, ‘16651’, and ‘24795’.

## Supporting Information Legends

**S1_Materials.docx** Additional information on phenotype derivation, mediation analysis and testing for indirect effect, *F*_*SNP*_, simulations, Supplementary Tables S1-S5, and Supplementary Figures S1-S2.

